# A novel mutation in the nucleoporin NUP35 causes murine degenerative colonic smooth muscle myopathy

**DOI:** 10.1101/036582

**Authors:** Ian A. Parish, Lincon A. Stamp, Ayla May D. Lorenzo, Suzanne M. Fowler, Yovina Sontani, Lisa A. Miosge, Debbie R. Howard, Christopher C. Goodnow, Heather M. Young, John B. Furness

## Abstract

Chronic Intestinal Pseudo-Obstruction (CIPO) is a rare, but life-threatening, disease characterized by severe intestinal dysmotility. Histopathological studies of CIPO patients have identified several different mechanisms that appear to be responsible for the dysmotility, including defects in neurons, smooth muscle or interstitial cells of Cajal. Currently there are few mouse models of the various forms of CIPO. We generated a mouse with a point mutation in the RNA Recognition Motif of the *Nup35* gene, which encodes a component of the nuclear pore complex. *Nup35* mutants developed a severe megacolon and exhibited reduced lifespan. Histopathological examination revealed a degenerative myopathy that developed after birth and specifically affected smooth muscle in the colon; smooth muscle in the small bowel and the bladder were not affected. Furthermore, no defects were found in enteric neurons or interstitial cells of Cajal. *Nup35* mice are likely to be a valuable model for the sub-type of CIPO characterized by degenerative myopathy. Our study also raises the possibility that *Nup35* polymorphisms could contribute to some cases of CIPO.

**Significance Statement:** Chronic Intestinal Pseudo-Obstruction (CIPO) is a disabling bowel disorder in which the symptoms resemble those caused by mechanical obstruction, but no physical obstruction is present. Some patients with CIPO have defects in intestinal neurons while in other CIPO patients the muscle cells in the bowel wall appear to degenerate, but the underlying cause of these defects is unknown in most CIPO patients. We generated a mouse that has a mutation in *Nup35*, which encodes a component of the pores found within the membrane of the cell nucleus. The mutant mice developed intestinal obstruction, which we showed was due to degeneration of the muscle cells in the colon. This mouse is likely to provide new insights into some forms of CIPO.

## Introduction

Chronic Intestinal Pseudo-Obstruction (CIPO) is a rare and debilitating condition in which deficiencies in intestinal peristalsis mimic a mechanical obstruction (1). CIPO can be caused by defects in the enteric nervous system (neuropathy), interstitial cells of Cajal and/or intestinal smooth muscle (myopathy) (1-5). While CIPO can occur secondary to other clinical complications, or downstream of environmental factors, certain forms of CIPO have a heritable genetic component (1, 6). For example, mutations in genes such as *SOX10, FLNA, L1CAM, RAD21, TYMP, ACGT2* and *POLG* have all been linked to CIPO pathology (3, 7-12). Nevertheless, the genetic cause of a large portion of heritable CIPO cases remains unknown (3).

Mouse models have previously been used to interrogate the role of genetic pathways in CIPO pathology (13). While mouse models have implicated the disruption of a number of genetic pathways in intestinal neuropathy, there is a paucity of models that mimic clinical intestinal smooth muscle myopathy. We report a novel mouse model of CIPO associated with a mutation in the RNA recognition motif (RRM) of the nucleoporin, NUP35. NUP35 (also called NUP53) is part of the multi-
protein nuclear pore complex (NPC), which is composed of multiple copies of ∼30 different nucleoporin proteins forming a complex greater than 40MDa in size (14). The NPC plays a critical role in allowing diffusion of small molecules and controlled movement of larger factors in and out of the nucleus. Proper NPC formation is also important for normal nuclear morphology. NUP35 is indispensable for vertebrate NPC formation and nuclear integrity (15-19), and NUP35 dimerisation via its RRM is critical for its function (19, 20).

Here we characterize the colonic changes associated with the NUP35 mutant CIPO mouse model, and show that the mice have colon-specific smooth muscle myopathy without detectable deficits in enteric neurons and with persistence of interstitial cells of Cajal. These data suggest that the NUP35 mutant mouse represents a novel model of CIPO associated with degenerative myopathy.

## Results

### Nup35^M92L/F^’l92l∼ mutant mice exhibit mortality associated with megacolon

As part of a larger ethylnitrosourea (ENU) mutagenesis screen for mutant mice with immune phenotypes, mice with a F192L mutation in the nucleoporin NUP35 were identified and isolated from the Australian Phenomics Facility PhenomeBank, and the mutation was bred to homozygosity. The crystal structure for the NUP35 RRM dimer has been solved (20), and F192, which is highly conserved (Fig. 1A), lies on the alpha helix of the dimer interaction interface (Fig. 1B). A substitution of phenylalanine for leucine, as occurs in the F192L mutation, will delete an aromatic ring that is buried within the hydrophobic core of the RRM (Fig. 1B) and is predicted to be highly damaging (Polyphen = 1, SIFT = 0). This mutation will thus likely disrupt the RRM interaction interface and thereby interfere with NUP35 dimerisation and function.

**Figure 1.**
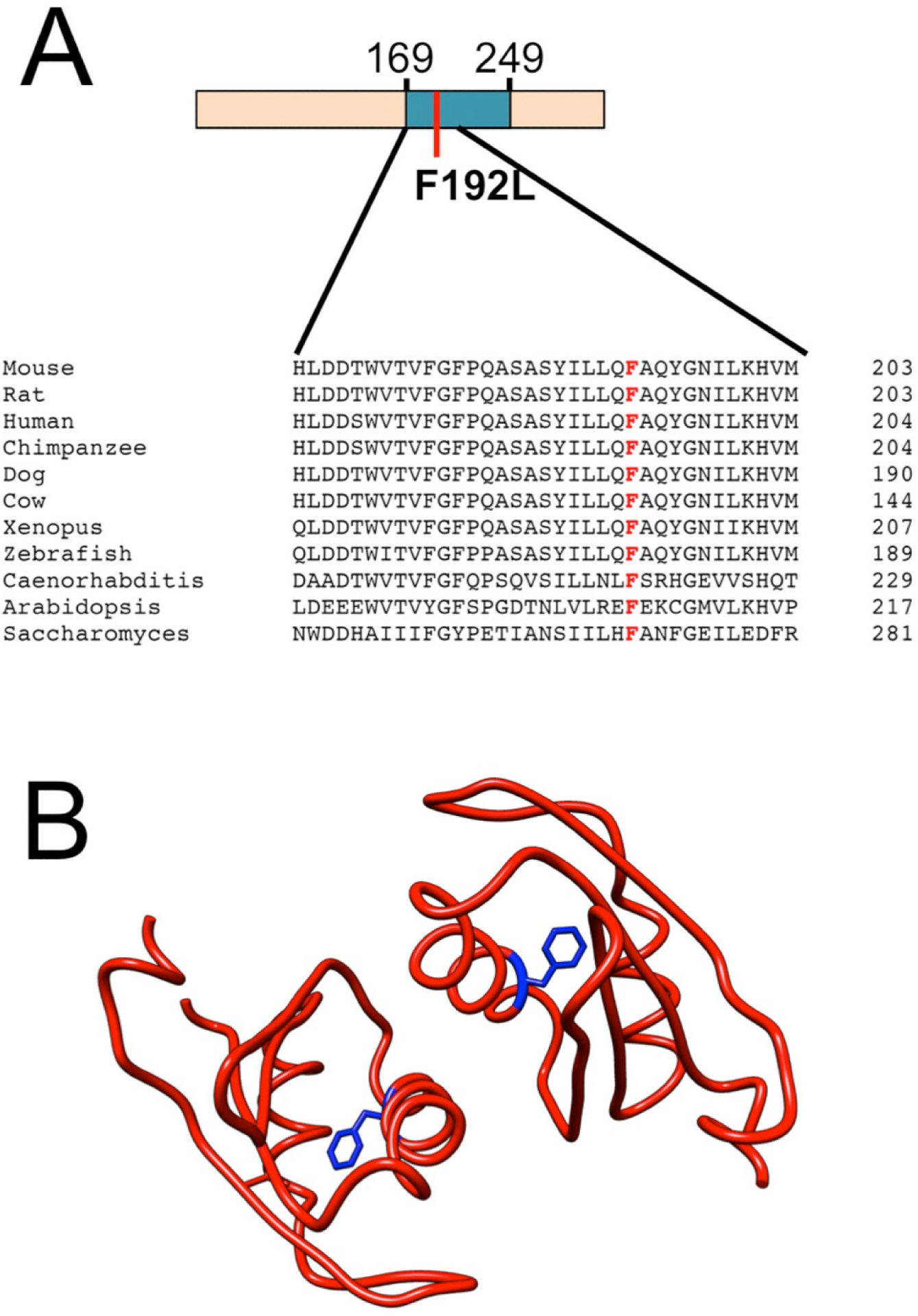
Conservation and structural positioning of the NUP35 F192 residue. **(A)** Sequence alignment showing NUP35 RRM amino acid conservation between species. Coloured schematic of mouse NUP35 protein shows the RRM in blue (which spans amino acids 169-249), with the location of the F192L mutated residue indicated by a red line. The F192 residue is also highlighted in the sequence alignment in red. **(B)** NUP35 RRM dimer crystal structure with F192 illustrated in blue and the orientation of the aromatic side chain shown.

Given the essential role of NUP35 in NPC formation and nuclear integrity (15-19), damaging mutations in this protein could be anticipated to cause embryonic lethality. However, while homozygous mutant mice were born below the expected 25% ratio, viable *Nup35*^F192L/F192L^ homozygous mutant mice were still obtained in appreciable numbers (Table 1). Homozygous mutant mice failed to display a detectable steady-state immune phenotype, however mortality was observed within this strain, with 50% of mice failing to survive beyond 55 days of age and no mice surviving beyond 120 days of age (Fig. 2A). These high mortality rates were not observed in *Nup35*^F192L/+^ heterozygous mutant mice (Fig. 2A). Post-mortem examination of mutant mice revealed substantial megacolon (Fig. 2B), with no obvious superficial pathology noted in the other organs examined (lung, liver, heart, bladder, spleen, thymus, kidney). The small intestine appeared normal, however faecal impaction was observed within the colon without any obvious physical obstruction. Thus, *Nup35*^F192L/F192L^ mutant mice develop symptoms consistent with human CIPO that appear to cause mortality.

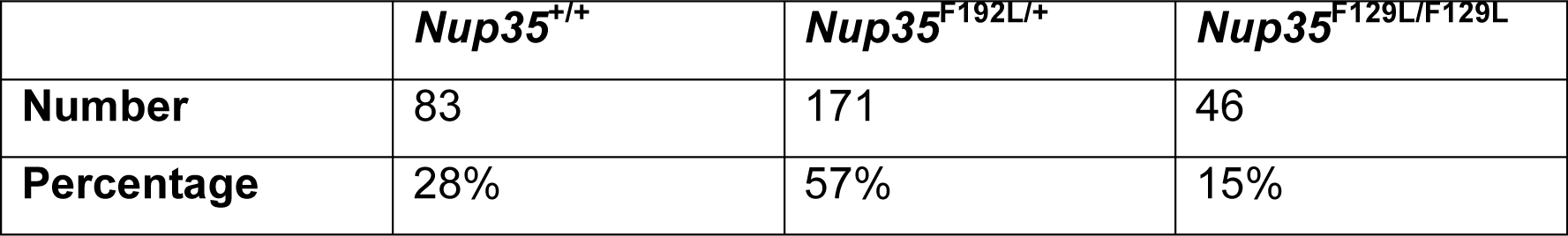
Partial embryonic lethality within *Nup35* mutant mice.

**Figure 2.**
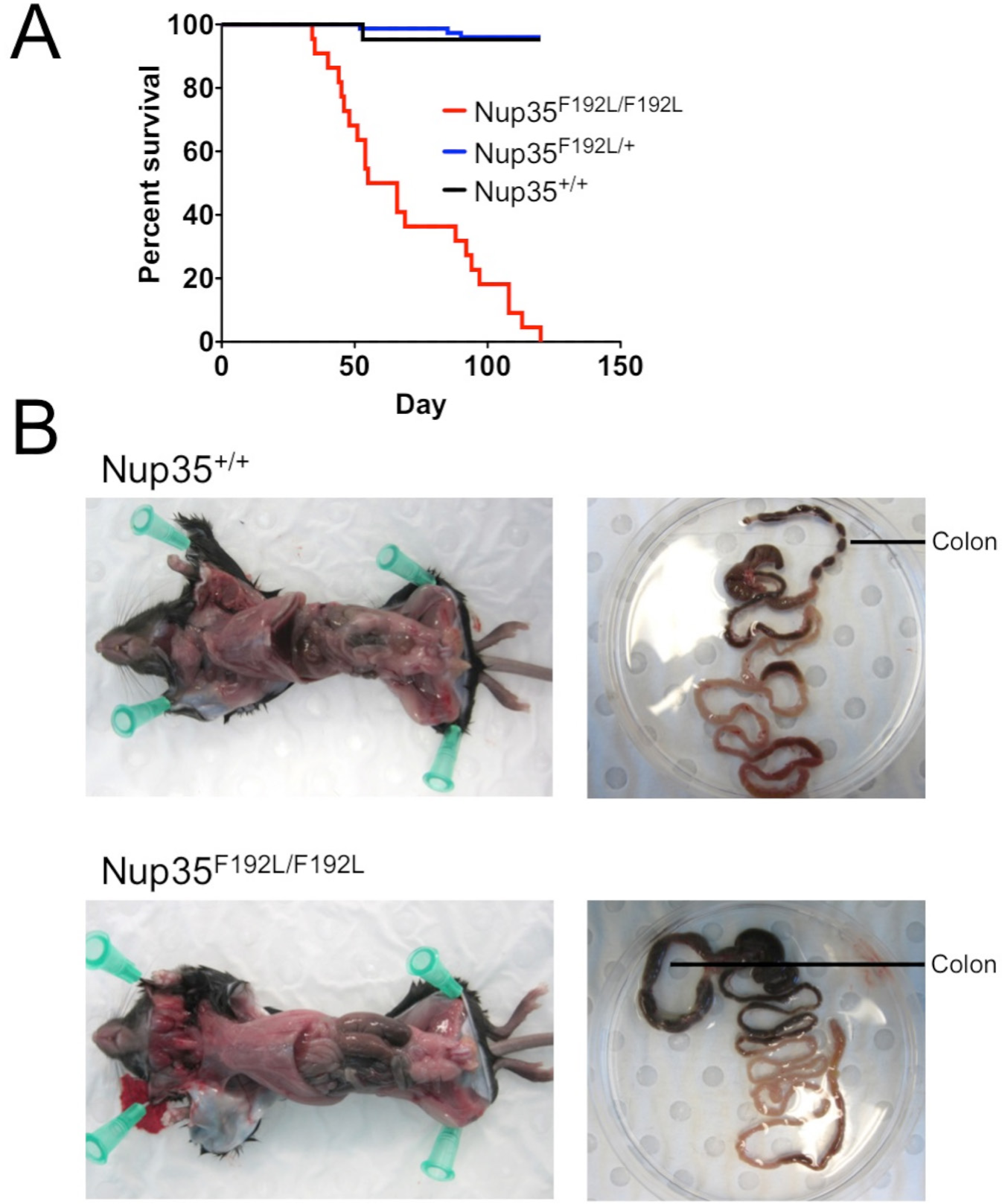
Mortality and megacolon within *Nup35*^F192L/F192L^ mutant mice. **(A)** Survival of *Nup35*^+/+^ (black line, n=21), *Nup35*^F192L/+^ (blue line, n=76) and *Nup35*^F192L/F192L^ (red line, n=26) mice. “Days” on the x-axis indicates post-natal days of age. Mortality was defined as mice that died without cause or were euthanised due to unknown illness. **(B)** Postmortem pictures illustrating megacolon within *Nup35*^F192L/F192L^ mice and wild-type *Nup35*^+/+^ littermates, showing colon size within the body cavity (left panels) or within the isolated intestines, with the location of the colon indicated (right panels).

ENU randomly introduces point mutations into the genome, and each strain may carry a number of passenger mutations that could be responsible for any observed phenotype. Based on exome sequencing data of the pedigree from which the *Nup35*^F192L/F192L^ mutant mice were derived, two other exome mutations existed on Chromosome 2 within this pedigree that could be loosely linked to the *Nup35* locus and may potentially contribute to the observed phenotype. These mutant alleles (in the genes *Cst10* and *Fbn1*) were subsequently found to be present within the *Nup35* mutant mouse colony. To confirm that these alleles did not contribute to the observed phenotype, a *Nup35* mutant mouse sub-line was established that lacked these two mutations, and mortality was still observed with comparable histopathology (see next section) to the parental mouse line (data not shown). Thus, the *Nup35* F192L mutation appears to be the causative mutation for the observed CIPO phenotype.

### Nup35^F192L/F192L^ mutant mice display degenerative mid- and distal colonic smooth muscle loss

CIPO can be caused by neuropathy, myopathy and/or defects in interstitial cells of Cajal, so we next employed histopathology and immunohistochemistry to determine the cause of bowel obstruction in *Nup35* mutants. Transverse sections of the ileum, and proximal, mid- and distal colon of 6-8 week old *Nup35*^F192L/F192L^ (n = 5), *Nup35*^F192L/+^ (n = 2) and *Nup35*^+/+^ (n = 6) littermates were stained with haematoxylin and eosin or Masson’s trichrome. No differences were observed in the ileum of *Nup35*^F192L/F192L^ mice compared to *Nup35*^F192L/+^ or *Nup35*^+/+^ mice. However, the distal and mid-colon of homozygous *Nup35*^F192L/F192L^ mice exhibited substantial loss of muscle cells and replacement by connective tissue in the muscularis externa, but not the muscularis mucosae (Fig. 3B,C,E,F). In some areas, no muscle cells were observed in the longitudinal and circular muscle layers, although the layering was still defined by connective tissue. The nuclei of smooth muscle cells in the circular muscle layer of the colon of *Nup35*^F192L/+^ and *Nup35*^+/+^ mice were very uniform in appearance (Fig. 3G). In contrast, the nuclei of the remaining muscle cells in *Nup35*^F192L/F192L^ mice were very variable; some had chromatin condensations (Fig. 3H), while others were indistinguishable from those in control mice. There were also patches of thick adherent mucus, and increased numbers of surface goblet cells that appeared to displace surface enterocytes in the mid- and distal colon. There was also thickening of the submucosa of *Nup35*^F192L/F192L^ mice, with greater numbers of macrophages being present.

**Figure 3.**
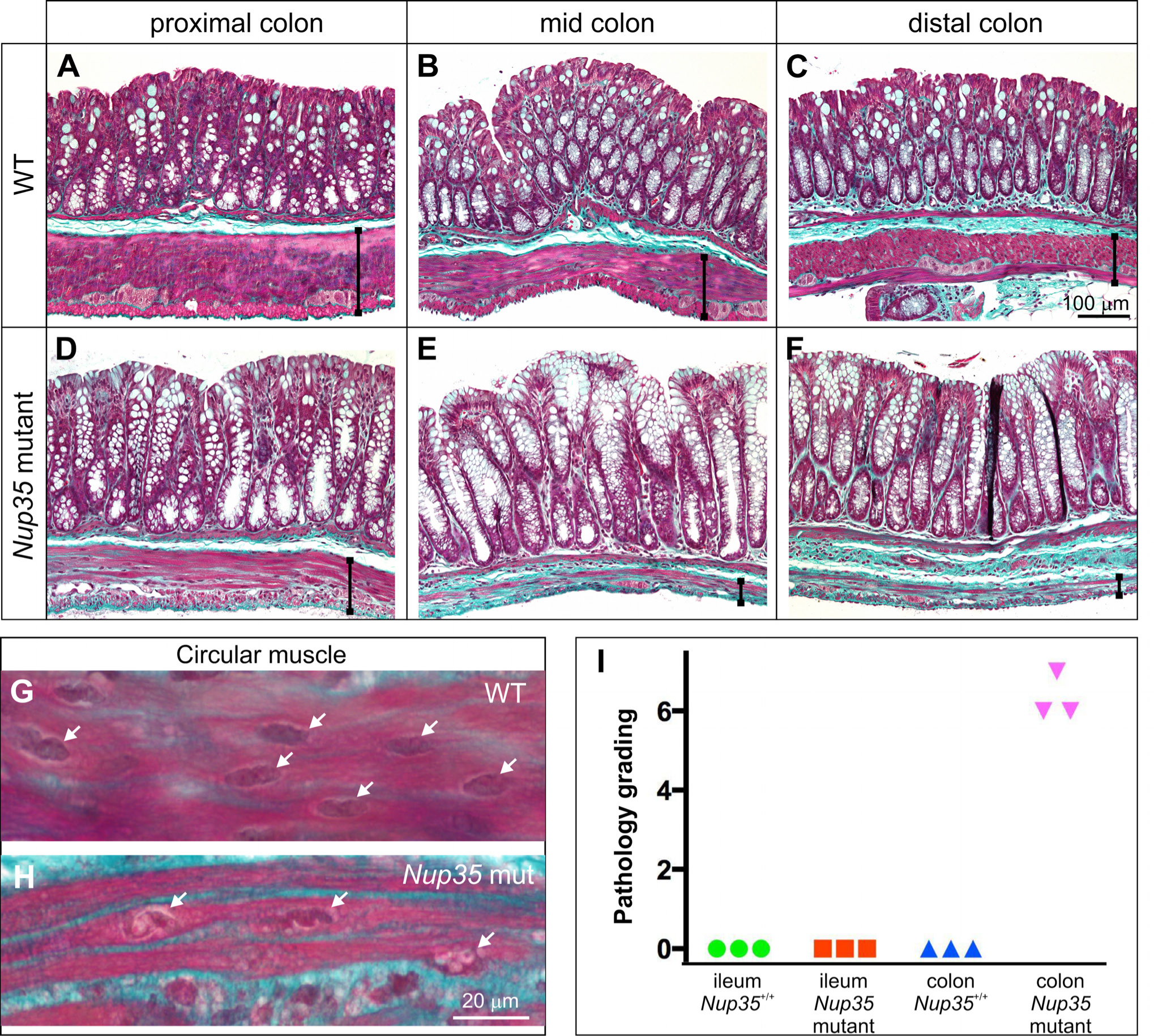
Smooth muscle loss from the external muscle layers of the colon of *Nup35* mutant mice. **A-F:** Masson’s trichrome staining of the proximal, mid- and distal colon from 6-8 week old wildtype (WT, **A-C)** and *Nup35*^F192L/F192L^ **(D-F)** mice. There is a dramatic loss of muscle cells from the circular and longitudinal muscle layers, and replacement by connective tissue (green), in the mid- and distal colon of the mutant mice. There is a less dramatic loss of muscle cells in the proximal colon. The width of the external muscle layers are indicated by vertical black lines. Note greater numbers of surface goblet cells **(E, F)** and thickened submucosa **(F)**. **G,H:** High magnification images of circular muscle cells. While the nuclei (*arrows*) of WT mice are uniform in appearance **(G)**, many nuclei (*arrows*) in the remaining smooth muscle cells in the mutant had chromatin condensations **(H). I:** Pathological grading of distal colon and ileum tissue from 3 randomly chosen *Nup35*^+/+^ and *Nup35*^F192L/F192L^ mice (0 is normal and 7 is severely affected), which included assessments of the longitudinal and circular muscle layers, the submucosa and mucosa (see Supplementary Tables 1 and 2). While the colons of all mutants were graded as severely affected, the ileum was graded normal.

The proximal colon of *Nup35*^F192L/F192L^ mice displayed only a minor myopathy (Fig. 3A,D), but like more distal regions, there was also a thick adherent mucus and more surface goblet cells. The histological characteristics of distal colon and ileum of *Nup35*^+/+^ and *Nup35*^F192L/F192L^ mice were quantified by a blind observer using 7 point scales (0 is normal and 7 is severely affected) that included appraisals of the longitudinal and circular muscle layers, the submucosa and mucosa (Supplementary Tables 1 and 2). The ileums of *Nup35*^F192L/F192L^ mice were graded normal, whereas the distal colons of *Nup35*^F192L/F192L^ mice were graded as severely affected (Fig. 3I). The bladder of homozygous NUP35 mutants appeared normal, with no detectable changes in the smooth muscle cells (data not shown). Striated muscle within the pelvic floor also appeared normal (data not shown), suggesting that the myopathy was highly restricted to the external smooth muscle within the colon.

To determine if the colonic myopathy is congenital or degenerative, sections of colon from newborn *Nup35*^F192L/F192L^ (n = 5), *Nup35*^F192L/+^ (n = 3) and *Nup35*^+/+^ (n = 3) littermates were stained with haematoxylin and eosin or Masson’s trichrome. The external muscle layers of *Nup35*^F192L/F192L^ newborn mice were similar in appearance to that of littermates, and no histological differences were detected between newborn homozygous mutant, heterozygous and wild-type mice, including the appearance of nuclei in the external muscle layers (Fig. 4).

**Figure 4.**
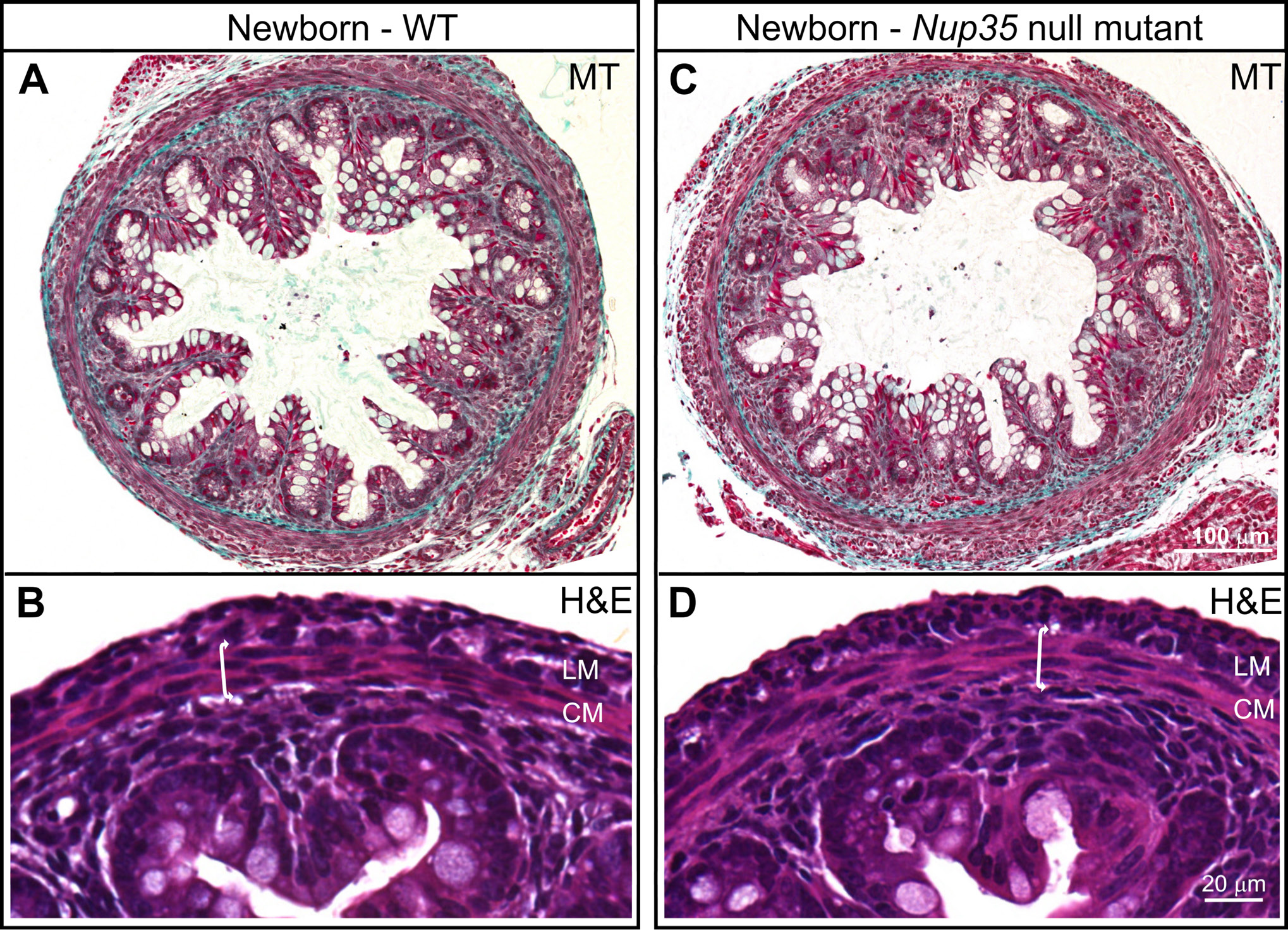
Newborn *Nup35*^F192L/F192L^ mutant colon does not exhibit any histological defects. Masson’s trichrome (MT, **A, C**) and haemotoxylin and eosin (H&E, **B, D**) staining of transverse sections of distal colon from newborn wildtype (WT, **A,B**) and *Nup35*^F192L/F192L^ (mutant, **C,D**) mice. There are no detectable differences between the longitudinal muscle layer (LM) and circular muscle layer (CM, and double arrow) between mutants and wild-type littermates.

Like smooth muscle cells, interstitial cells of Cajal (ICC), macrophages in the muscularis layers and enteric neurons play an essential role in gut motility (21, 22). For example, congenital absence of enteric neurons from the distal bowel results in bowel obstruction and megacolon (23, 24). Wholemount preparations of the external muscle layers of the ileum and colon from 6-8 week old mice of *Nup35*^F192L/F192L^, *Nup35*^F192L/+^ and *Nup35*^+/+^ littermates were processed for immunohistochemistry using antisera to the pan-neuronal marker, Tuj1, the enteric neuron sub-type marker, nNOS, the ICC marker, Kit, and the macrophage marker, F4/80. Enteric neurons (Fig. 5A and D) and the ICC network (Fig. 5B and E) were present along the entire bowel, although ICC in the circular muscle layer of the colon of the mutants had fewer fine processes than those in wild-type mice (Fig. 5E). There were many more macrophages in the external muscle and associated with myenteric ganglia, particularly in the mid- and proximal colon of homozygous NUP35 mutants compared to heterozygote and wild-type littermates (Fig. 5C and F). Collectively, these data show that the megacolon in NUP35 homozygous mutant mice appears to be due to a degenerative myopathy of the colon.

**Figure 5.**
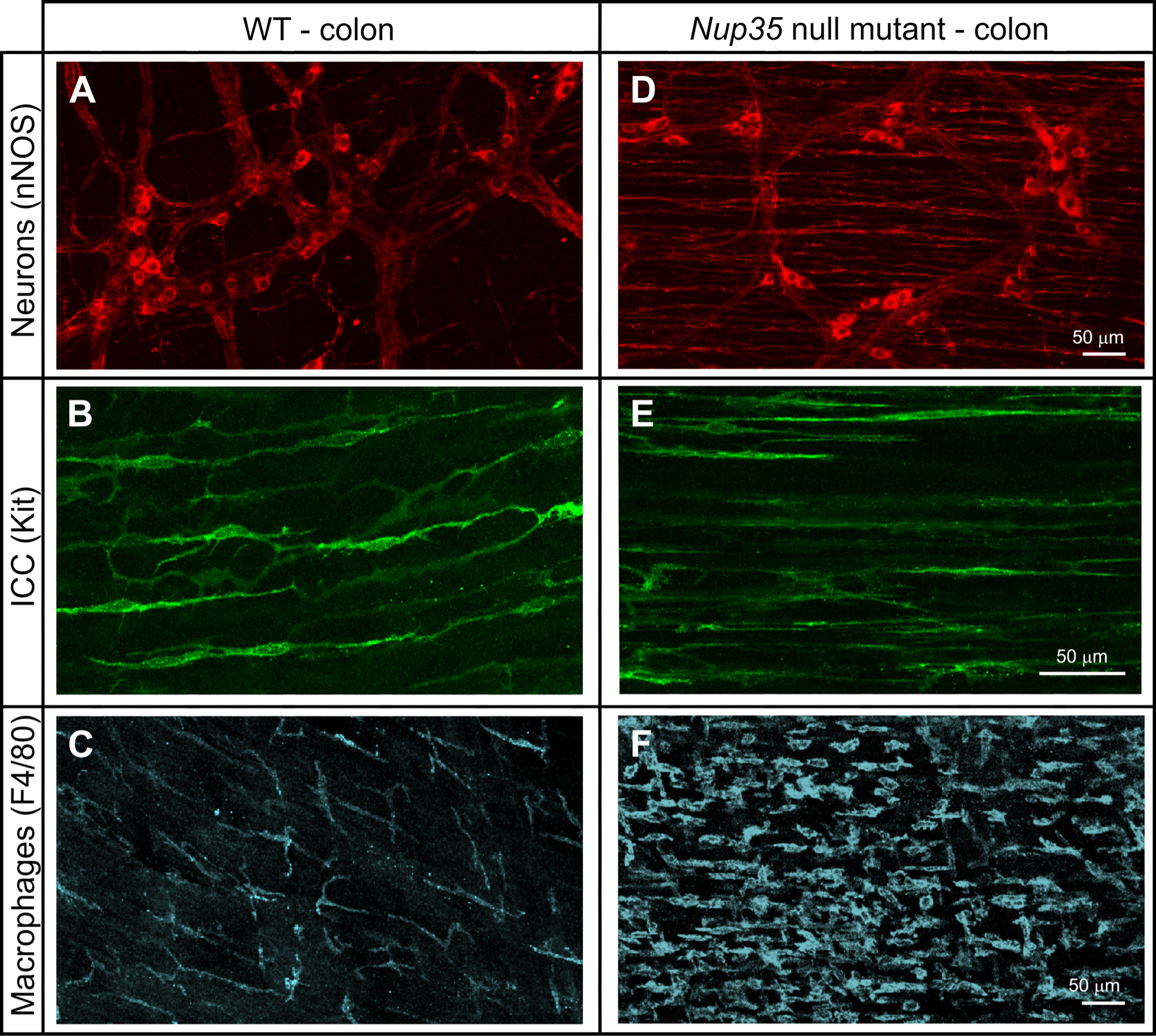
Neurons and interstitial cells of Cajal (ICC) are present along the colon of *Nup35*^F192L/F192L^ mutant mice. Wholemount preparations of external muscle and myenteric plexus from the distal colon (**A, D**) and mid-colon (**B-F**) of 6-8 week old WT (**A-C**) and *Nup35* mutant (**D-F**) mice following immunostaining for the enteric neuron sub-type marker, nNOS (**A, D**), the ICC marker, Kit (**B, E**) and the macrophage marker, F4/80 (**C, F**). **A, D:** nNOS+ myenteric neurons in the distal colon. **B, E:** Although circular smooth muscle cells have been largely replaced by connective tissue, Kit+ ICC are present in the circular muscle layer of mutants. However, ICC in the mutants have simpler morphology and fewer processes than ICC in the circular muscle of WT mice. **C, F:** F4/80+ macrophages are more rounded in shape and are more abundant in the circular muscle and myenteric region of *Nup35* mutants compared to WT littermates.

## Discussion

We describe a novel mouse model of colonic myopathy and CIPO associated with a mutation in the nucleoporin NUP35. The phenotype associated with the NUP35 mutation is surprising at two levels. First, NUP35 is reported to be indispensable for NPC formation and nuclear integrity (15-19), and it is essential for *C. elegans* embryonic development (18), so it is surprising that viable mutant mice were obtained. Second, the phenotype observed is highly selective, with smooth muscle myopathy only observed in the muscularis externa of the colon despite no evidence of tissue-specific NUP35 expression within existing expression databases such as BioGPS (http://biogps.org/), and the Human Protein Atlas (http://www.proteinatlas.org/). Overall, these data reveal an unexpected link between the nuclear pore complex and CIPO, and suggest that *Nup35* polymorphisms could contribute to disease.

The phenotype within the *Nup35* mutant mouse appears distinct to other gene deficient myopathy models and, to our knowledge, this is the first gene-deficient mouse model demonstrating spontaneous degenerative smooth muscle myopathy and associated fibrosis. Previous models of myopathy associated with CIPO-like disease have involved inducible deletion of the transcriptional regulator, serum response factor (SRF), from smooth muscle (25, 26), or deletion of the smooth muscle-restricted factor Smoothelin A (27). In both cases, the CIPO phenotype was associated with loss of smooth muscle cell contractility rather than active loss of smooth muscle cells; SRF deletion failed to affect smooth muscle cell numbers, while Smoothelin A loss triggered smooth muscle cell hypertrophy. The large phenotypic differences between the *Nup35*^F192L/F192L^ mouse and the other models described above make it unlikely that *Nup35* deficiency causes myopathy by disrupting SRF or Smoothelin A, implicating a potentially novel pathway in the *Nup35*^F192L/F192L^ mouse phenotype.

As smooth muscle fibrosis was reported in a clinical case of Hollow Visceral Myopathy (a form of CIPO associated with degenerative myopathy) (28), the *Nup35*^F192L/F192L^ mutant mouse represents a potentially valuable model for human disease. Moreover, unlike the congenital neuropathy, Hirschsprung disease (29, 30), CIPO patients with degenerative myopathy are variable in age (3). The colonic myopathy in *Nup35* mutant mice was not apparent at birth and so is consistent with the degenerative myopathy seen in a sub-population of CIPO patients.

The histopathological characteristics of CIPO varies between patients and include neuropathy, loss of ICC as well as degenerative myopathy (3). Mutations linked to human CIPO are often associated with neuropathy rather than myopathy (eg. *SOX10* and *RAD21*) (3, 12), and so are unlikely to be related to the *Nup35* mutant phenotype. The mechanism by which NUP35 deficiency triggers smooth muscle myopathy thus remains unclear. Mutations in *FLNA* are associated with myopathy in X-linked CIPO (8), however such myopathy is associated with abnormal (additional) layering of the small intestinal muscularis propria rather than smooth muscle cell loss (11). Myopathy associated with fibrosis is observed in patients suffering from mitochondrial neurogastrointestinal encephalomyopathy, a multifactorial disease with CIPO symptoms (9). These patients bear mutations in *TYMP*, which leads to mitochondrial depletion from smooth muscle that likely causes myopathy (9). There is no reported association between either NUP35 and TYMP, or NUP35 and other mitochondrial components, however we cannot rule out that mitochondrial abnormalities contribute to myopathy in these mice.

One possible explanation for the phenotype is that nuclear abnormalities associated with NUP35 deficiency cause myopathy. Polymorphisms in *LMNA*, encoding LAMIN A, a structural factor that lines the nucleus, have been linked to striated muscle wasting and muscular dystrophy (31, 32). LAMIN A is also a relatively widely expressed protein, so the reasons for the tissue-selective effect of *LMNA* polymorphisms on striated muscle maintenance remain unclear. One theory is that the more fragile nucleus that results from LAMIN A loss is susceptible to rupture in muscle due to mechanical stress (31, 32). The subsequent loss of nuclear integrity is postulated to cause cell death. The NPC is known to associate with Lamins, and interestingly a direct association between NUP35 and LAMIN B has been reported (17). Furthermore, NUP35 depletion leads to nuclear abnormalities that closely resemble those seen in LAMIN A mutant cells (17). It is thus possible that NUP35 deficiencies cause disrupted nuclear morphology that triggers cell death in contractile smooth muscle cells. Consistent with this idea, we did observe some degree of abnormal nuclear morphology within the mutant smooth muscle cells, although these changes could simply be due to ongoing apoptosis. However, some degree of tissue-specificity is still required for this mechanism to operate, as altered nuclear morphology should also cause striated muscle loss, which we failed to observe in *Nup35* mutant mice. Furthermore, cell loss was only observed in a small subset of smooth muscle cells within the body (colonic smooth muscle cells), suggesting a complex highly tissue-
specific effect.

In some cases, CIPO is associated with inflammation, predominantly of the enteric ganglia, which exhibit inflammatory neuropathy (1). Neither inflammation nor enteric nervous system damage was observed in tissue from the *Nup35* mutant mice. However, we did observe increased numbers of macrophages in the affected external muscle and the adjacent submucosa, which is possibly a response to smooth muscle cell degeneration. Changes in the morphology of ICC may also be secondary to the myopathy and the loss of muscle cells. ICC form gap junctions with smooth muscle cells (33), and the simpler morphology of ICC in *Nup35* mutants is likely to be due to reduced contacts with smooth muscle cells. The accumulation of gut contents, pressure on the gut wall, and failed propulsion may induce the increased numbers of surface goblet cells and the adherent mucus that was observed.

As NUP35 does not exhibit clear tissue-specific expression, it is unclear why we observed such a selective phenotype in *Nup35* mutant mice. There is emerging evidence of heterogeneity within NPC composition between tissues, which is highlighted by the tissue-specific pathologies associated with loss of other nucleoporins (14). It is thus possible that the NPC present within colonic smooth muscle is particularly susceptible to NUP35 depletion, leading to selective myopathy within this cell type. Regardless of the mechanism, the Nup35^F192L/F192L^ mutant mouse phenotype has revealed a novel pathway involved in smooth muscle myopathy that may contribute to human CIPO.

## Methods

### Mice

*Nup35*^F192L/F192L^ mice were isolated from the Australian Phenomics Facility PhenomeBank at the Australian National University. All animals used in this study were cared for and used in accordance with protocols approved by the Australian National University Animal Experimentation Ethics Committee and the current guidelines from the Australian Code of Practice for the Care and Use of Animals for Scientific Purposes.

### NUP35 sequence alignment and protein modeling

NUP35 sequences were isolated from UniProt (http://www.uniprot.org/) and aligned using Clustal W (34). Protein structure visualisation was performed with the UCSF Chimera package, which was developed by the Resource for Biocomputing, Visualization, and Informatics at the University of California, San Francisco (supported by NIGMS P41-GM103311) (35).

### Immunohistochemistry and histopathology

Mice were killed by cervical dislocation, and the bladder, colon and small intestine removed and fixed overnight in formalin. Tissue for H&E and Masson’s trichrome staining was processed as described previously (36). Tissue from 3 randomly chosen sections of ileum and distal colon were graded by a blinded observer using a X40 objective lens and 7 point graded scales (Supplementary Tables 1 and 2).

Tissue for wholemount immunohistochemistry was opened along the mesenteric border and processed for immunohistochemistry as described previously (37) using the following primary antisera: rabbit anti-Kit (1:100, Calbiochem; (38)), mouse anti-Tuj1 (1:2000, Covance), sheep anti-nNOS (1:2000; (39)) and rat anti-F4/80 (1:50; (40)). Secondary antisera (all from Jackson ImmunoResearch) were: donkey anti-rat Alexa488 (1:100), donkey anti-rabbit Alexa647 (1:400), donkey anti-mouse Alexa594 (1:200) and donkey anti-sheep Alexa594 (1:100). Wholemount preparations were imaged on a Zeiss Pascal confocal microscope.

### Data analysis

Graphing and data analysis was conducted using Prism Software (GraphPad).

## Acknowledgements

We would like to kindly thank the Australian Phenomics Facility staff for assistance with animal work, and the Histopathology and Organ Pathology Service of the Australian Phenomics Network. This work was supported by an NHMRC CJ Martin Fellowship (I.A.P.), NHMRC Program Grant 1016953 and NIH grant AI100627 (C.C.G.), NHMRC Senior Research Fellowship APP1002506 (H.M.Y.) and NHMRC Project Grants APP1079234 (H.M.Y., L.A.S.) and APP1005811 (J.B.F.).

